# RNA Immunoprecipitation and Sequencing of Isolated RNAs (RIP-SIR) Identifies Endogenous miRNA-Target Interactions

**DOI:** 10.1101/2022.08.22.504764

**Authors:** Julie Wells, Anuj Srivastava, Richard S. Maser, Rupesh Kesharwani, Vishal Kumar Sarsani, Sandra L. Daigle, Ryan P. Lynch, Heidi J. Munger, Rosalinda Doty, Ruth L. Saxl, Carol J. Bult

## Abstract

MicroRNAs (miRNAs) are crucial post-transcriptional regulators of gene expression in species ranging from plants to mammals. Unraveling the complex networks of miRNA-mediated regulation of gene expression requires the ability to identify the mRNA targets of individual miRNAs in native cellular contexts. Here we describe RIP-SIR, an unbiased genome-wide method for identifying interactions between endogenous miRNAs and their targets in almost any tissue or cell type.

---

miRNAs (miRNAs) regulate gene expression post-transcriptionally by direct base-pairing interactions between nucleotides 2-7 of the miRNA with complementary nucleotides within target mRNAs, known as miRNA response elements (MREs), resulting in degradation or inhibited translation of bound transcripts ^1,2^. Identifying the targets of individual miRNAs is complicated by the existence of non-canonical miRNA binding sites and miRNA isomers as well as the influence of cellular context on miRNA-MRE interactions ^3,4^. Current computational and experimental techniques for identifying miRNA targets cannot fully recapitulate the biological complexity of target selection. For example, it is difficult for miRNA target prediction algorithms to incorporate the effects of spatial and temporal co-expression patterns, stoichiometry and threshold levels of miRNAs and target mRNAs; interactions between target mRNAs and RNA binding proteins; and the multiple sources for input transcript annotation which can vary significantly among 3’UTR sequence^5,6^. Furthermore, most miRNA target-prediction algorithms limit their searches for MREs to the 3’UTR of transcripts ^7^ despite increasing experimental evidence for their location also within 5’UTRs, exons, introns and intergenic regions of the genome ^8^.

Experimental methods for identifying the targets of individual miRNAs typically require expression of exogenous, tagged miRNAs which may perturb native stoichiometry, are limited to the study of cultured cells and cannot be used to monitor miRNA-mediated regulation during processes such as differentiation or disease progression. Even methods such as Argonaute <u>Hi</u>g<u>h</u>-<u>T</u>hroughput <u>S</u>equencing of RNAs isolated by <u>C</u>ross<u>L</u>inking and <u>I</u>mmuno<u>P</u>recipitation (Ago HITS-CLIP) and <u>C</u>ross<u>L</u>inking <u>A</u>nd <u>S</u>equencing of <u>H</u>ybrids (CLASH), cannot match individual miRNAs to their targets or rely upon very low efficiency intramolecular ligation, respectively ^9,10^. To overcome the limitations of current methods for miRNA target identification, we developed <u>R</u>NA <u>I</u>mmuno<u>P</u>recipitation and <u>S</u>equencing of <u>I</u>solated <u>R</u>NAs (RIP-SIR) and demonstrate that RIP-SIR is capable of identifying which miRNAs are bound to individual transcripts in lung tissues from both healthy mice and those bearing late-stage pulmonary adenocarcinomas.

RIP-SIR comprises two phases, which together identify the individual miRNAs interacting with transcripts of interest (Fig. 1a). In the first phase, referred to as RIP-seq, miRNAs and their bound miRNA binding sites are collectively identified. In the second phase, referred to as SIR, only the miRNAs bound to an individual binding site are identified. To demonstrate the applicability of RIP-SIR to studying miRNA binding during disease progression, we employed a mouse model of pulmonary adenocarcinoma. Following exposure to an adenovirus expressing Cre recombinase (adenoCre), early-stage (ET) animals developed predominantly adenomas ^11^ while late-stage (LT) animals developed predominantly adenocarcinomas ^12^ (Supplemental Fig. 1). As negative controls, animals from both models were either untreated (Un) or infected with an adenovirus that does not express Cre recombinase (adeno) and are referred to as early-stage normal (EN) or late-stage normal (LN) respectively. Lungs were treated with formaldehyde to form crosslinks between miRNAs, target mRNAs and individual proteins of bound RNA induced silencing complexes (RISCs). Crosslinked miRNA-mRNA-RISC complexes were immunoprecipitated by the addition of an antibody against AGO2, one component of RISC, and magnetic beads conjugated to protein A (Fig. 1a). Additional samples received either an antibody against IgG, as a negative control, or no antibody (labeled as 10% input), as a positive control (Fig. 1b). Protein-RNA complexes isolated from each sample were washed and eluted from the magnetic beads, crosslinks were reversed and protein components were digested.

**Fig. 1.**
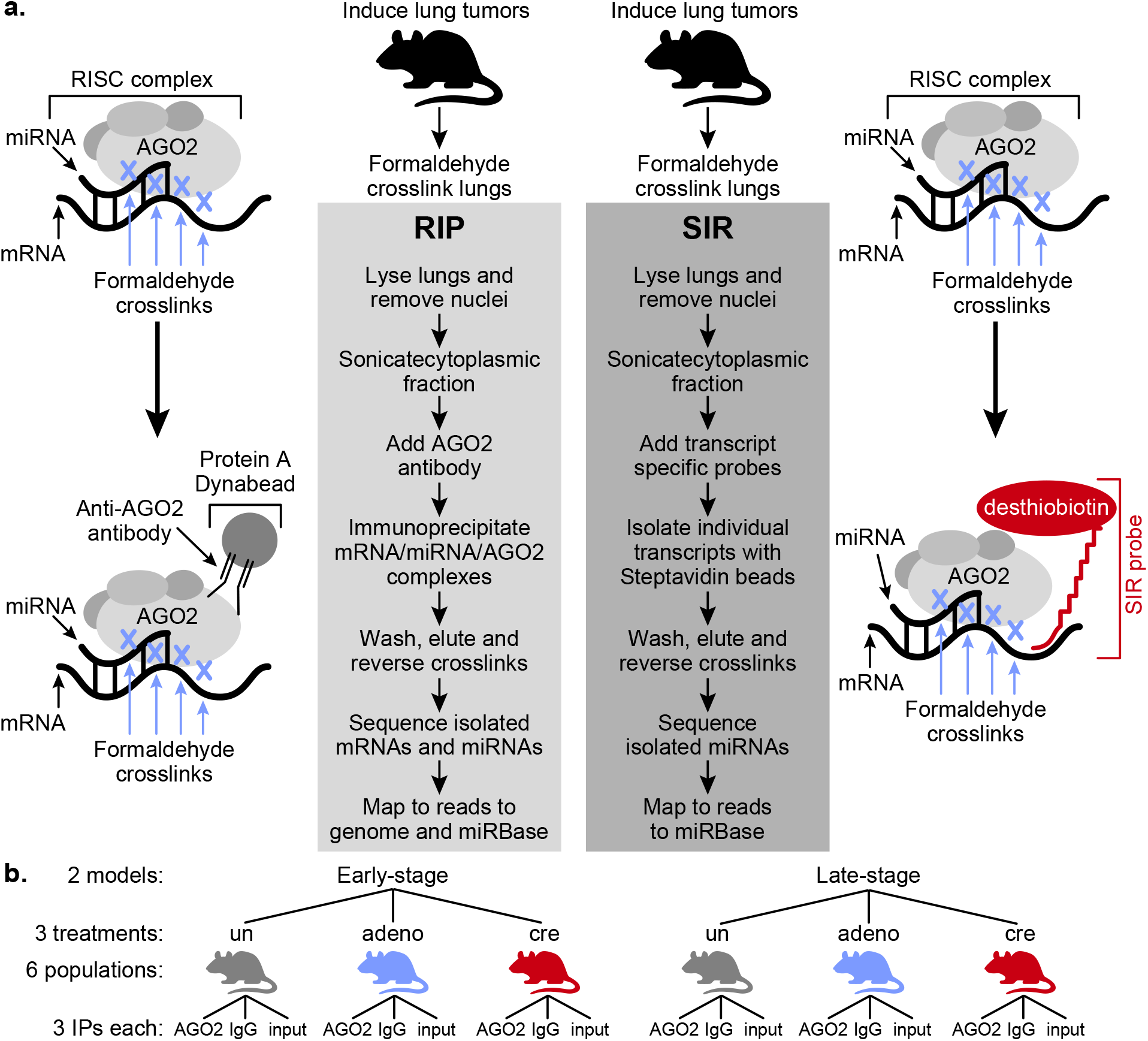
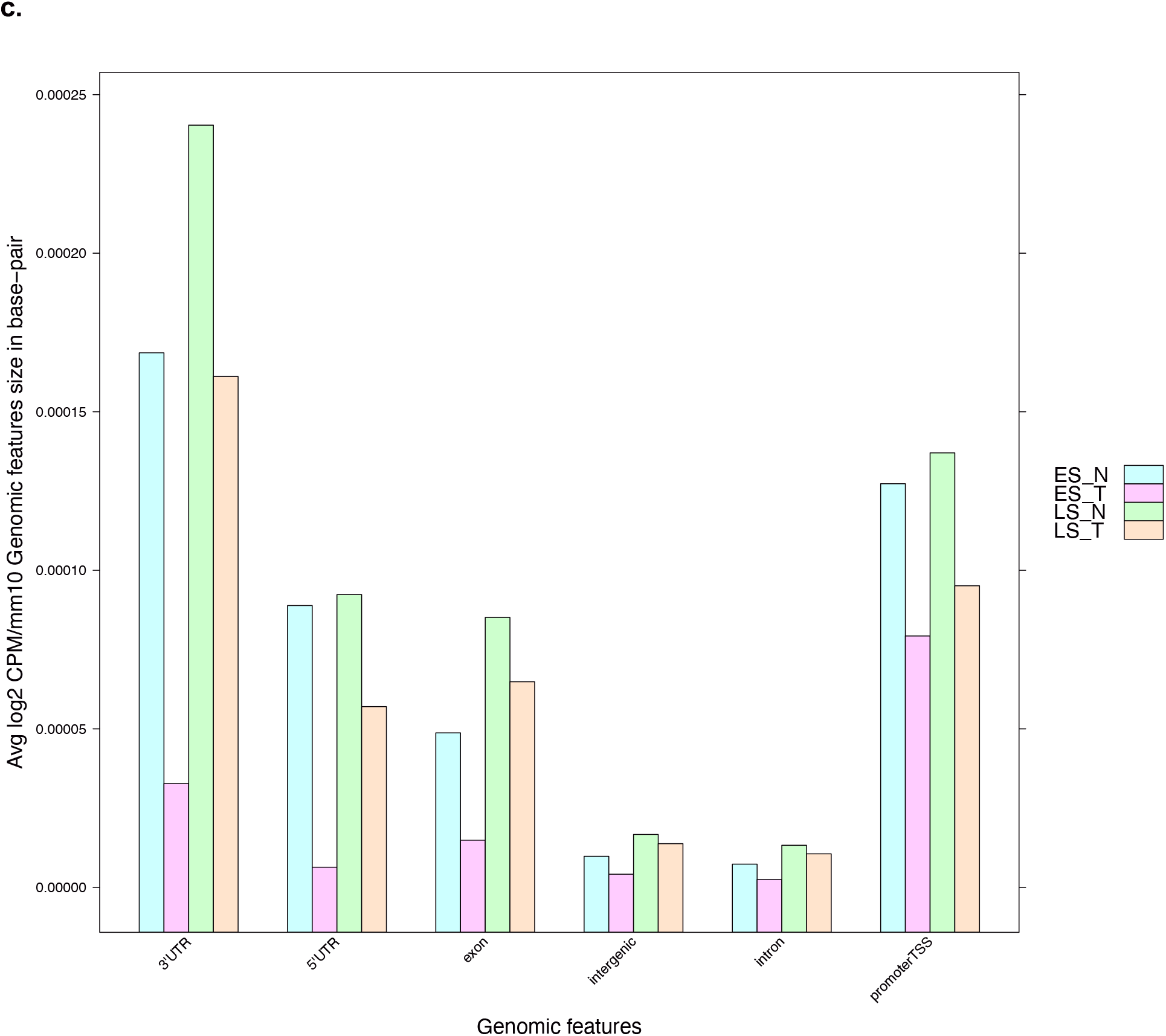
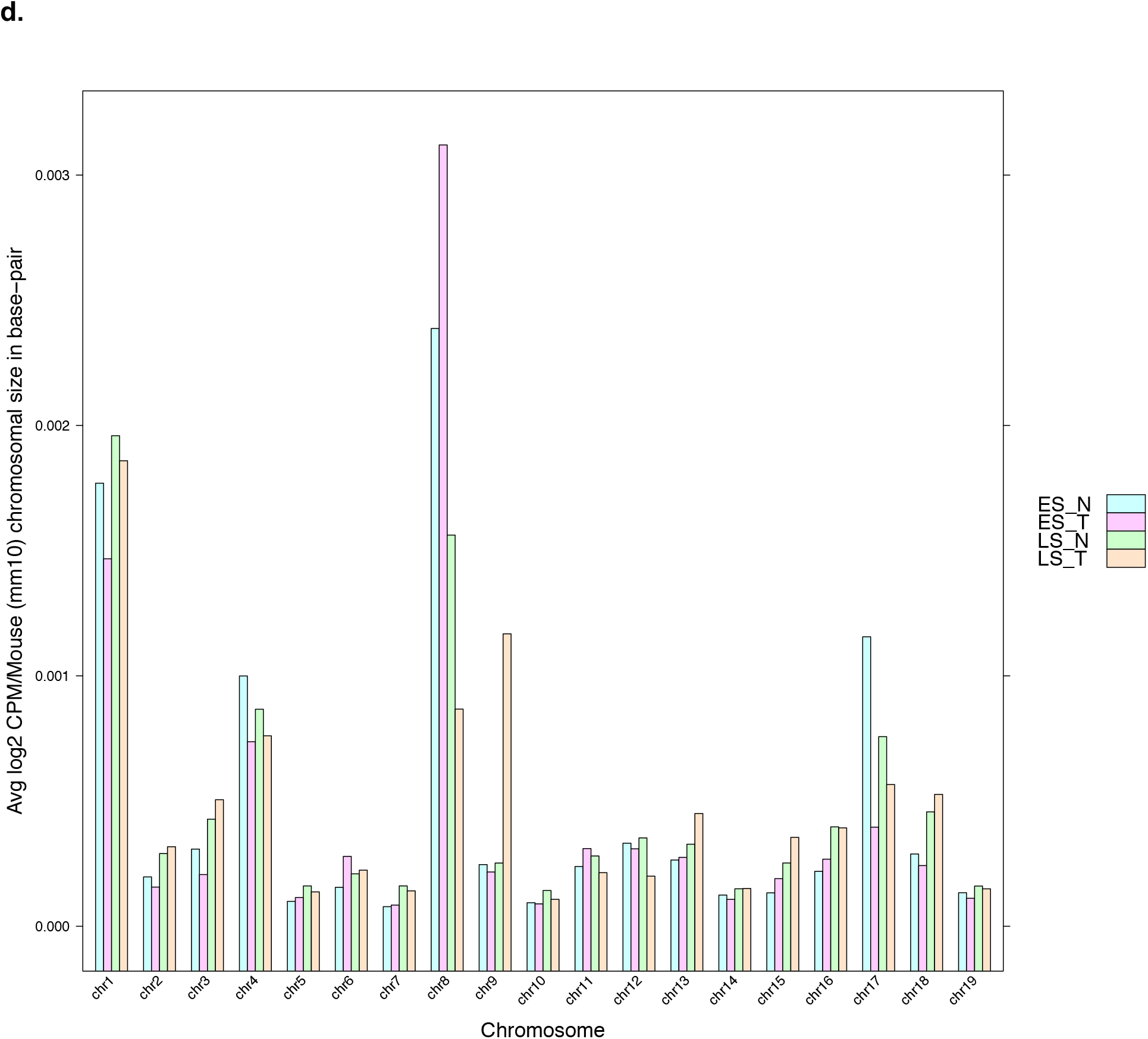
RIP-SIR Identifies miRNA targets in a mouse model of pulmonary adenocarcinoma. **a**, Schematic representation of the RIP-SIR assay. The panel on the left represents the RIP-seq portion of the assay and identifies all bound miRNAs and all bound target sequences. The panel on the right represents the SIR portion of the assay and identifies which miRNAs are bound to an individual transcript. **b**, Schematic representation of the mice used for RIP-seq assays. Mice from both the early-stage (K) and late-stage (KP) tumor models were divided among three different treatment groups: uninfected (un), animals infected with adenovirus (adeno) and animals infected with adenovirus expressing Cre recombinase (cre). Crosslinked chromatin from three mice from each treatment group was pooled and immunoprecipitated with an antibody against argonaute 2 (AGO2), an isotype control antibody (IgG) or was not immunoprecipitated (input). **c**, The genomic locations of all mapped RIP-seq mRNA reads were determined and annotated with a genomic feature, 3’UTR, 5’UTR, exon, intergenic, intron or promoter transcriptional start site (TSS). Total read counts are plotted for each genomic feature as count per million, normalized by the length of each feature in the GRCm38 genome assembly build. **d**, The location of each mapped RIP-seq mRNA read is mapped to a single chromosome and read counts are displayed as counts per million, normalized to the size of each chromosome in the GRCm38 genome assembly build.

High-throughput sequencing libraries were generated from the RNAs isolated from each sample and gel purified. Two fractions were collected and individually sequenced from each library: the first fraction contained cDNAs generated from RNA templates of ~20 nts to ~80 nts in length representing miRNAs and small RNA species and the second fraction contained cDNAs of between ~81 nts to ~500 nts representing bound target mRNAs (Supplemental Fig. 2). mRNA sequencing reads were mapped to the GRCm38 genome assembly build while miRNA reads were mapped using miRBase v21. We sequenced an average of 3272415, 1415094, 3092071 and 4554125 RIP miRNA reads per group for EN, ET, LN and LT, respectively. After quality control, ~7.5 to 11% of these reads were successfully mapped to miRbase v21 (Supplemental Table 1). Similarly, we sequenced 3506908, 1863381, 2908163 and 4242283 mRNA reads per group for EN, ET, LN and LT, respectively, of which 2.5 to 4.5% were successfully mapped to the GRCm38 genome assembly build (Supplemental Table 1). We further performed differential expression analysis to identify miRNAs and target mRNAs with inversely correlated expression patterns between tumor (ET and LT) and normal (EN and LN) samples (Supplemental Table 3). Mapping of the mRNA reads to the mouse genome identified from 1 to 8 MREs per transcript (Supplemental Table 4). In agreement with recent findings of MREs outside of target transcript 3’UTRs, a large portion of our Ago2 binding sites mapped to exons, introns, intergenic regions and transcriptional start sites (Fig. 1c). Putative MREs were mapped to every chromosome, reflecting the large portion of the mouse genome thought to be regulated by miRNAs (Fig. 1d).

From among the 200 target transcripts with the highest number of reads identified by RIP-seq analysis, we selected four transcripts for SIR analysis: protein phosphatase 1 regulatory subunit 16b (*Ppp1r16b*; MGI:2151841), PR domain zinc finger 9 (*Prdm9*; MGI:2384854), Notch 2 (MGI: 97364) and high mobility group 2 (*Hmga2*; MGI:101761), as a positive control. The 3’UTR of *Hmga2* transcripts contain 8 putative binding sites for *let-7*, six of which are functional for miRNA-mediated repression of Hmga2 expression in luciferase reporter assays ^13^. We designed five probes for SIR, each containing 25nts of RNA separated from a desthiobiotin molecule by a 108 atom tethering arm, mimicking the design of Pitch probes ^14^. Every other RNA residue, beginning with the second, contained a 2’fluoro (2’F) linkage modification. The RNA portion of the *Hmga2* SIR probe was complementary to a region of the *Hmga2* transcript near several of the previously mapped *let-7* miRNA binding sites and the MRE we mapped by RIP-seq (Fig. 2a). A second probe, referred to as “scrambled”, contained the same nucleotides as the HMGA2 probe in random order as a negative control. The four remaining probes were complementary to regions of selected transcripts (*Prdm9, Ppp1r16b* or *Notch2*) adjacent to an MRE mapped by RIP-seq analysis (Fig. 2a, Supplemental Table 1). We generated probes near each of mapped MREs within the Notch2 transcript to determine if SIR could detect binding of different miRNAs to different MREs within the same transcript.

**Fig. 2.**
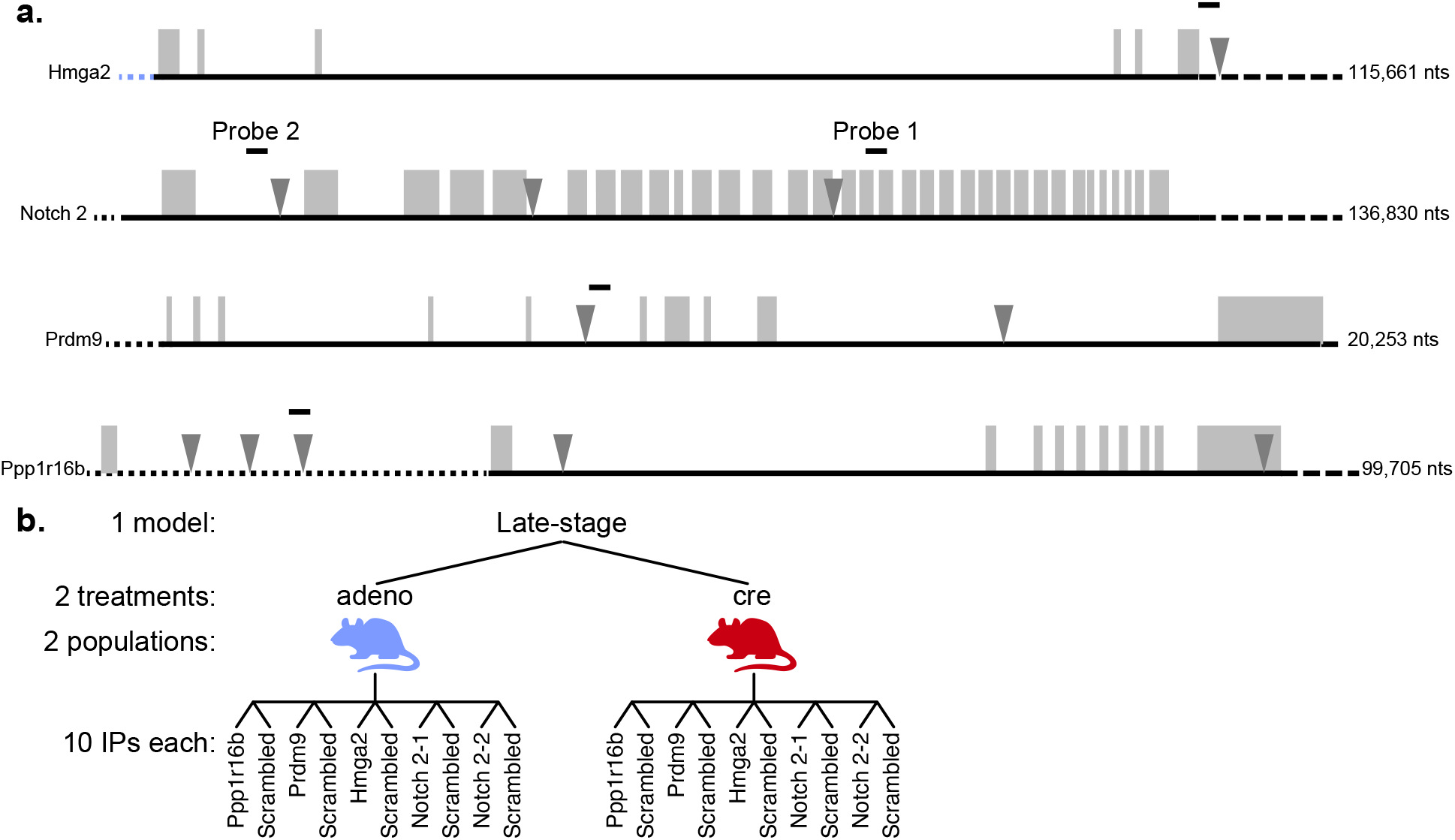
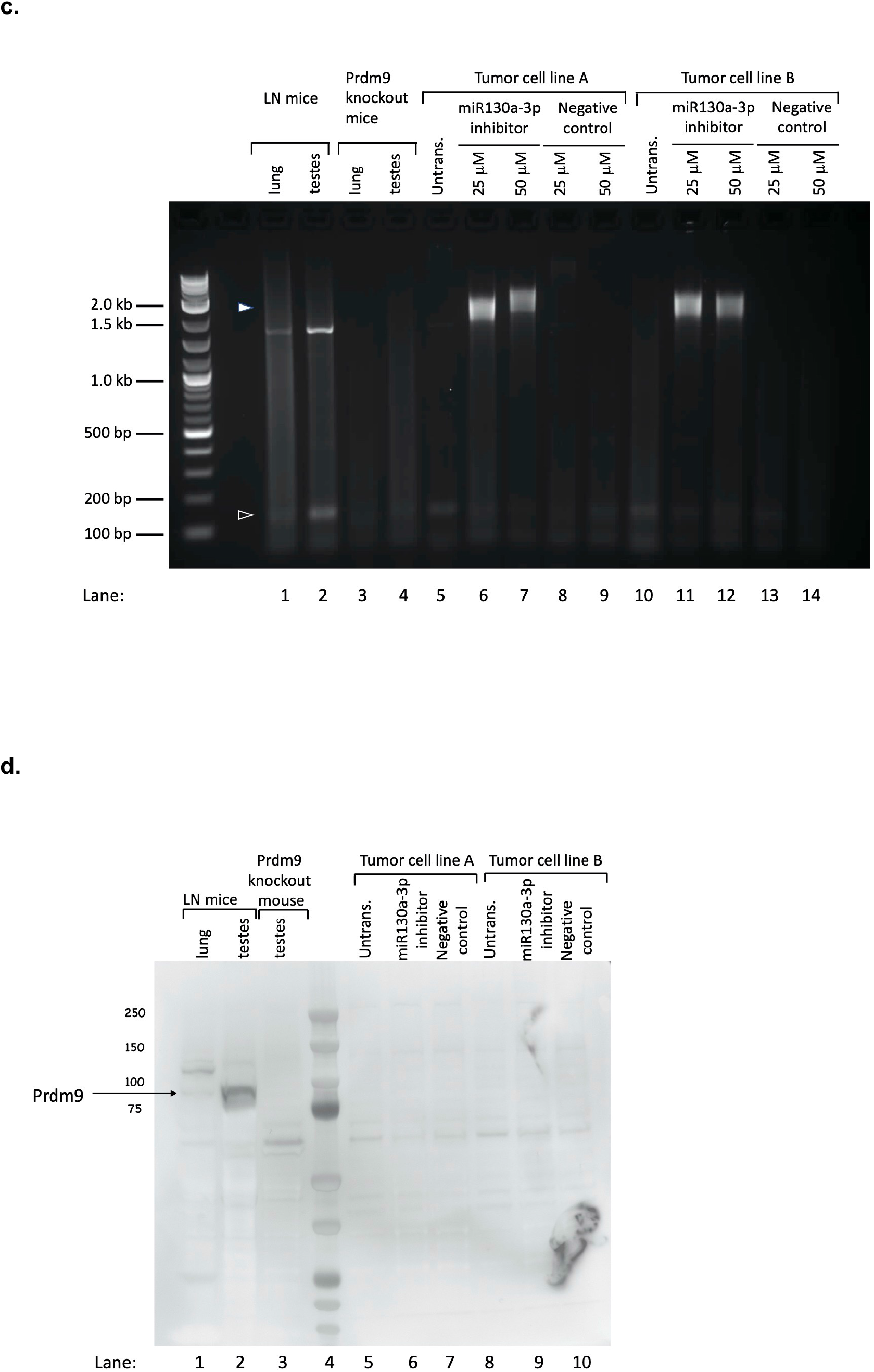
SIR identifies the miRNAs bound to individual transcripts. **a**, Schematic representations of the positive control transcript (Hmga2) and three experimental transcripts (Notch 2, Prdm9 and Ppp1r16b) chosen for SIR analysis. For each transcript, dotted lines represent 5’UTR sequences, solid lines represent coding sequence regions and dashed lines represent 3’UTR sequences. Shaded rectangles represent exons, inverted triangles indicate the locations of MREs mapped by RIP-seq and heavy, short lines above each transcript show the locations of SIR probes. The sizes of the transcripts are indicated at right. Diagram is not drawn dto scale. **b**, Schematic representation of the mice used for SIR analysis. Mice from the late-stage tumor model were divided between two treatment groups: those infected with adenovirus (adeno) and those infected with adenovirus expressing Cre recombinase (cre). Crosslinked chromatin from four mice was pooled for SIR analysis with either one of five transcript specific probes or a scrambled probe as a negative control. **c**, PCR analysis of Prdm9 expression. RNA was isolated from lung or testes tissues of KP mice (lanes 1–2) from lung or testes tissues of Prdm9 knock-out mice (lanes 3–4), KP tumor cell line A (lanes 5–9) or B (lanes 10–14) and reverse transcribed into cDNA. Tumor cells were untransfected (lanes 5, 10), transfected with 25 (lanes 6,10) or 50 (lanes 7, 11) micromolar inhibitor against mmu-miR130a-3p or 25 (lanes 8,12) or 50 (lanes 9, 13) micromolar non-specific inhibitor. **d**, Western blot analysis of Prdm9 protein expression shows low level expression in normal mouse lung tissue (lane 1) and higher expression in normal testes tissue (lane 2) but not in testes tissue from a Prdm9 knock out mouse (lane 3). Prdm9 protein levels are very low in KP mouse lung tumor cell lines A and B (lanes 5, 8) and expression levels do not change following transfection with 50 micromolar of either an inhibitor against mmu-miR130a-3p (lanes 6,9) or a negative control (lanes 7, 10).

For SIR analysis, lungs from late-stage pulmonary adenocarcinoma animals infected with either adenoCre or adeno were formaldehyde crosslinked (Fig. 2b). Crosslinked miRNA-target mRNA complexes were isolated by the addition of a SIR probe followed by streptavidin beads. After extensive washing, complexes were eluted, crosslinks were reversed and the liberated miRNAs were sequenced. As positive controls, sequencing libraries were also generated from 10% of the starting input from each sample. High-throughput sequencing analysis and mapping of the sequencing reads to the mm10 genome build identified numerous miRNAs bound to each mRNA in both late-stage tumors and normal lung tissues (Table 1). Our SIR analysis identified binding of predominantly *miR-181a-1-3p*, not *let-7*, to the *Hmga2* transcript. We detected one or two predominant miRNAs bound to every putative MRE, plus lesser amounts of additional miRNAs (Table1). Comparison of SIR results obtained from tumors and normal tissues indicates a high degree of overlap in the identity, but not always the abundance, of miRNAs bound to each MRE in the two tissue types. We demonstrated MREs identified by RIP-SIR and located outside of 3’UTRs are functional by generating tumor cell lines from LT animals and measuring Prdm9 expression levels before and after transfection with a microRNA hairpin inhibitor against mmu-miR-130-3a. Inhibition of *miR-130a-3p* resulted in the appearance of an approximately 2 kb *Prdm9* PCR product (Fig. 2c) but did not affect expression of PRDM9 protein (Fig. 2d). Sequencing analysis of this PCR product showed that it was generated by the retention of intron 5 within *Prdm9* transcripts, suggesting a role for miRNAs in regulating intron retention.

**Table 1.**
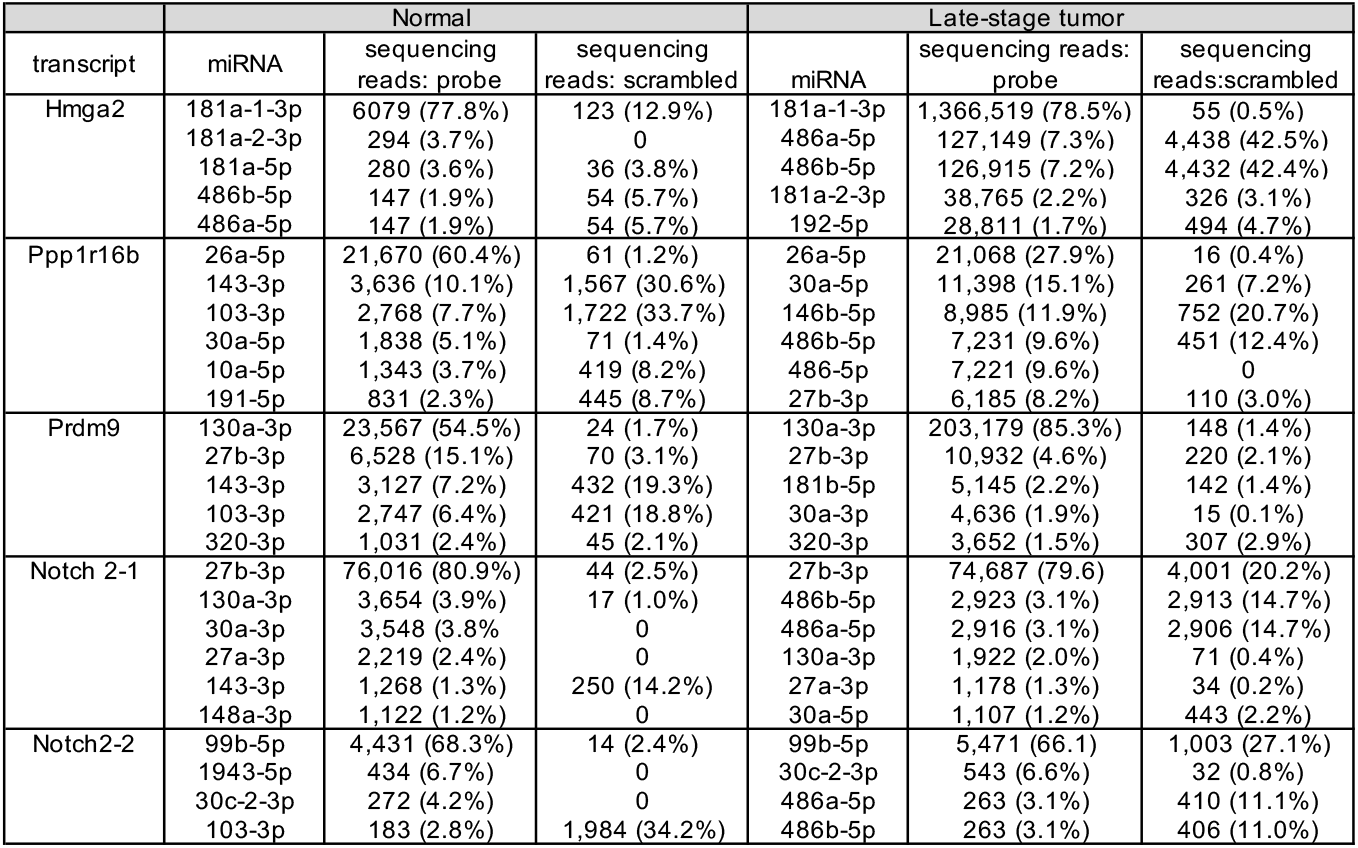
Mapped read counts of miRNAs bound to individual MREs in normal lungs and late-stage pulmonary adenocarcinoma

Deciphering how miRNAs regulate expression of target genes remains a complex problem. Here, we have demonstrated that RIP-SIR identifies the miRNAs bound to individual MREs under native conditions, even in complex tissues during disease progression. RIP-SIR may also prove valuable in identifying the targets of other short RNA regulators, such as isomirs, piwi-interacting RNAs and tRNA-derived fragments, whose abundance is dependent upon sex, origin, race, tissue type and disease state ^4^.

## Methods

### Mice

For RIP-SIR analysis, we obtained two strains of mice: B6.129S4-Kras^tm4Tyj^/J (stock #008179), referred to as K, and B6.129P2-Trp53^tm1Brn^/J (stock #008462), referred to as P. Both K and P mice were obtained from The Jackson Laboratory (Bar Harbor, ME) and were interbred to create mice that were heterozygous for the Kras^G12D^ allele and homozygous for the floxed Trp53 allele, referred to as KP. Male K and KP mice underwent one of three different experimental treatments at 6-7 weeks of age. The first treatment group of each mouse strain, referred to as early-stage tumor (ET) or late-stage tumor (LT), respectively, received intranasal administration of 5×10^6^ PFU of adenovirusCre (Viral Vector Core, University of Iowa, Iowa City, IA). The second group received 5×10^6^ PFU of adenovirus (Viral Vector Core, University of Iowa, Iowa City, IA) while the third group was not infected. Collectively, K animals from the second and third groups are referred to early-stage normal (EN) and KP animals from the second and third groups are referred to late-stage normal (LN). Animals in the first two treatment groups were anesthetized with intraperitoneal injections of tribromoethanol (400 mg/kg body weight) prior to viral instillation. Animals in all three treatment groups were harvested 16 weeks later, at 22 weeks of age. The left lung lobe of each animal was stained with hemotoxylin and eosin, embedded, sectioned and examined by microscopy for the expected histology. ET mice developed numerous, small lesions that were predominantly adenomas and a few adenocarcinomas. LT mice developed a greater number of larger lesions that were almost entirely adenocarcinomas. None of the EN or LN mice developed detectable lung lesions. All animal work was performed according to IACUC approved protocol #10111 at The Jackson Laboratory.

For analysis of Prdm9 expression, we collected lung and testes tissues from a homozygous, 6 week old, male B6;129P2-Prdm9^TM1YMAT/J^ mouse (Jackson Laboratory, Bar Harbor, ME stock #01071^15^.

### RNA immunoprecipitation (RIP)

At 22 weeks of age, animals in all treatment groups were euthanized by cervical dislocation and their lungs were perfused with 10 mls of cold 1X phosphate buffered saline (137mM NaCl, 2.7mM KCl, 10 mM Na2HPO4, 1.8mM KH2PO4 [PBS]) followed by 5 mls of 0.25% formaldehyde (Fisher Scientific Fair Lawn, NJ) diluted in 1X PBS. Lungs were removed and placed in 0.25% formaldehyde and rotated at room temperature for 15 minutes. Crosslinking reactions were stopped by the addition of glycine (Fisher Scientific Fair Lawn, NJ) to 0.125 M. Following dissection to remove unwanted tissues, crosslinked lungs were cut into small pieces using a razor blade then dissociated into single cell suspensions using Medicon filters (BD Biosciences San Jose, CA), washed with cold 1X PBS plus 1 mM phenylmethylsulfonyl fluoride (PMSF, Sigma-Aldrich St. Louis, MO) four times and pelleted by microcentrifugation. Cell pellets were resuspended in RIP lysis buffer (0.2% nonidet P40 substitute [Fluka/Sigma-Aldrich St. Louis, MO], 1 mM PMSF [Sigma-Aldrich St. Louis, MO] 0.01 mg/ml aprotinin [Sigma-Aldrich St. Louis, MO] 0.01 mg/ml leupeptin [Sigma-Aldrich St. Louis, MO] and 40 units/ml RNasin Plus [Promega, Madison, WI] pH 8.0) and incubated on ice for 20 minutes. Samples were processed with a dounce homogenizer and a small aliquot was visually examined with a microscope to ensure cell membranes had ruptured but nuclei remained intact. Samples were centrifuged to pellet nuclei and the cytoplasmic fractions were transferred to clean tubes. Cytoplasmic fractions were sonicated using a Bioruptor (Diagenode Denville, NJ) for 5 pulses of 30 seconds on high followed by 30 seconds off, centrifuged and supernatants were again transferred to clean tubes and stored at −80°C. For immunoprecipitations, cytoplasmic fractions collected from 3 mice of the same genotype and treatment group were pooled for each RIP reaction and precleared with 25 microliters of Dynabeads protein A resuspended in RIP lysis buffer (Invitrogen Life Technologies, Grand Island, NY) and rotated at 4°C for 30 minutes. Supernatants were divided among three tubes: one tube contained 10% of the supernatant fraction (input sample) and did not receive antibody while 45% of the supernatant was transferred to each of two tubes. One tube received 1 microgram of rabbit Ago2 antibody (Abcam #ab32381 Cambridge, MA) while the second received 1 microgram of rabbit IgG (Sigma-Aldrich, St. Louis, MO) as a negative control and rotated overnight at 4°C. Crosslinked antibody/protein/RNA complexes were isolated by the addition of 25 microliters of Dynabeads resuspended in RIP lysis buffer and rotation at 4°C for 15 minutes followed by microcentrifugation. Pellets were washed two times with 1.4 mls of RIP Wash-1000 buffer (I M NaCl, 1% nonidet P40 substitute, 0.1% sodium deoxycholate [Sigma-Aldrich, St. Louis, MO], 0.05% sodium dodecyl sulfate [SDS Fair Lawn, NJ] 1mM PMSF in 1X PBS) and four times with RIP Wash buffer (1% nonidet P40 substitute, 0.1% sodium deoxycholate, 0.05% SDS and 1 mM PMSF in 1X PBS). Complexes were eluted by the addition of 100 microliters of crosslink reversal buffer 50 mM Tris-Cl, pH 7.5, 5 mM EDTA, 1% SDS and 10 mM dithiothreitol [Fisher Scientific, Fair Lawn, NJ] and incubation at 70°C for 45-60 minutes with shaking. Samples were centrifuged, supernatants were transferred to clean tubes and RNAs were purified using mirVana PARIS kits (ThermoFisher, Waltham, MA) following manufacturer’s instructions. In the final purification step, RNA was eluted by the addition of RNase-free water and evaporated to dryness before library preparation. To estimate the efficiency of our fractionation, we examined the number of reads from all 18 of our 10% input samples that mapped to either β-actin or GAPDH. Both β-actin and GAPDH are enriched in soluble nuclear cell fractions^16^. Collectively, 5 reads mapped to β-actin (range: 0-3 per sample) and 3 reads mapped to GAPDH (range: 0-2 reads per sample).

### Sequencing of isolated RNAs (SIR)

Similar to RIP as described above, animals in all treatment groups were euthanized by cervical dislocation at 22 weeks of age and their lungs were perfused with 10 mls of cold 1X phosphate buffered saline, followed by 5 mls of 1.0% formaldehyde (Fisher Scientific Fair Lawn, NJ) diluted in 1X PBS. Lungs were removed and placed in 1.0% formaldehyde and rotated at room temperature for 15 minutes. Crosslinking reactions were stopped by the addition of glycine (Fisher Scientific Fair Lawn, NJ) to 0.125 M. Following dissection to remove unwanted tissues, crosslinked lungs were cut into small pieces using a razor blade then dissociated into single cell suspensions using Medicon filters (BD Biosciences San Jose, CA), washed with cold 1X PBS and 1 mM PMSF four times and pelleted by microcentrifugation. Cells from each animal were resuspended in 400 microliters of RIP lysis buffer and incubated on ice for 30 minutes. Following dounce homogenization, samples were centrifuged, supernatants were transferred to clean tubes and stored at −80°C.

Crosslinked supernatants were thawed and samples from 4 mice were combined for each SIR sample, for a total volume of ~1.5 mls per sample. One microliter of DNase I (Promega, Madison, WI), 2 microliters of RNasin Plus were added to each sample and samples were incubated at 37°C for 30 minutes. Ethyleneglycol-bis-(b-amino ethyl ether) N,N,N’,N’-tetraacetic acid (EGTA, Sigma-Aldrich St. Louis, MO) was added to a final concentration of 2.5 mM and samples were incubated at 37°C for an additional 15 minutes. 2.5X SIR Hyb buffer (50 mM HEPES, pH 7.5, 2 M NaCl, 4.75 M Urea, 1% SDS, 14.25 mM EDTA, 0.075% sodium deoxycholate and 12.5X Denhardts solution) was added to a final concentration of 1X and samples were sonicated using a Bioruptor for 4 cycles of 30 seconds on high and 30 seconds off. Fifty microliters of Ultralink Streptavidin (Pierce Thermo Scientific Rockford, IL), previously washed with 1 ml of 1X SIR Hyb buffer was added to each sample and samples were rotated at room temperature for 2 hours. Samples were centrifuged and divided among tubes to receive different SIR probes. One tube from each sample received a transcript specific probe while the other sample received a scrambled probe. Six microliters of 100 micromolar probe were added for every 1 ml of sample. Probes were synthesized by Fidelity Systems, Inc. (Gaithersburg, MD) and the sequences of each probe are listed below:

Hmga2: desthiobiotin-108 atom tethering arm-GtAaGaGtAcAgAgAaGaAtGgUcg
Ppp1r16b desthiobiotin-108 atom tethering arm-aCtGtCtGgAgCtUgUaGaGaGaGc
Prdm9: desthiobiotin-108 atom tethering arm-aAgGgCcCaCtGcAaUtAgGcAtUg
Notch2-1: desthiobiotin-108 atom tethering arm-aGtGcUcUgAtGaAcAcUgUgAcUg
Notch2-2: desthiobiotin-108 atom tethering arm-aGgUtGaGaUcCcAgGcGtGcAcAg
Scrambled: desthiobiotin-108 atom tethering arm-GgAcGtCaAgAgAtGaAtAgAgTa

Capitalized letters are 2’F-RNA residues and lowercase letters are DNA residues.

Samples were transferred to PCR strip tubes, 150 microliters per tube, and hybridized in a thermocycler for one cycle of 50°C for 20 minutes, 37°C for 10 minutes, 45°C for 60 minutes, 37°C for 30 minutes followed by a hold at 4°C. After hybridization, samples were transferred to 1.5 ml centrifuge tube and centrifuged at 16,000 x g for 15 minutes and supernatants were transferred to clean 2 ml tubes. Forty microliters of MyOne Streptavidin C1 beads (Invitrogen Life Technologies, Oslo, Norway), previously washed with 800 microliters of 1X SIR Hyb buffer, were added to each sample and mixed gently. Samples were rotated overnight at room temperature, placed on a magnetic stand and supernatants were transferred to clean tubes and stored at −80°C. The magnetic beads were washed 7 times with 1 ml of WB250 buffer pH 8.0 (250 mM NaCl, 10 mM HEPES, 2 mM ethylenediamine tetraacetic acid [EDTA], 1 mM EGTA, 0.2% SDS and 0.1% N-laurylsarcosine). Beads were carefully transferred to clean tubes before the last wash and resuspended in 100 microliters of elution buffer (12.5 mM biotin [Molecular Probes Life Technologies] in WB250 buffer) and incubated at room temperature with shaking for 60 minutes. The temperature was increased to 65°C and samples were incubated with shaking for an additional 10 minutes. Samples were immobilized on a magnetic stand, supernatants were transferred to clean tubes, one volume of 2X Crosslink reversal buffer (50 mm Tris-Cl, pH 7.5, 2 mM EDTA, 1% SDS and 10 mM DTT) was added. Samples were incubated at 70°C for 45 minutes then stored at −80°C^14,17,18^. The stored supernatants were thawed and subjected to a second round of SIR, beginning with the addition of SIR probes. Supernatants from samples that received a transcript specific probe in the first round received scrambled probe for the second round and vice versa. After the second round of SIR was completed, duplicate samples were combined and RNA was purified using the mirVana PARIS kit, following manufacturer’s instructions and eluting each sample in 100 microliters of nuclease-free water. Samples were evaporated to dryness and stored at −80°C.

### Tissue histology

The left lung lobe of each mouse was fixed in neutral buffered formalin (Labchem, Inc. Pittsburgh, PA) overnight at room temperature. Tissues were transferred to 70% ethanol and stored at room temperature. Fixed tissues embedded in paraffin, sectioned, stained with hemotoxylin and eosin and microscopically examined for the presence of tumors.

### Library construction and sequencing

Sequencing libraries were constructed starting with the entire pellet of RIP or SIR RNA, using TruSeq Small RNA kits from Illumina (San Diego, CA), following manufacturer’s instructions. Two different gel fragments were isolated from each gel of RIP samples. The first fragment, containing miRNAs, corresponded to a size of 145 bps-160 bps, including ligated adapters. The second fragment of each gel, containing immunoprecipitated mRNA targets, corresponded to a size of 160 bps-500 bps, including ligated adapters. Only one gel fragment, containing miRNAs corresponding to a size of 145 bps-160 bps, including ligated adapters, was isolated from each SIR sample. Libraries were checked for quality and concentration using the DNA High-Sensitivity LabChip assay (Agilent Technologies, Santa Clara, CA) and quantitative PCR (KAPA Biosystems, Wilmington, MA), according to the manufacturers’ instructions. RIP-seq libraries were pooled and sequenced on a MiSeq 50 base pair single-read flow cell using v2 reagents (Illumina, San Diego, CA). SIR libraries were also pooled and sequenced on either a HiSeq2500 high output 100 base pair paired-end flow cell using TruSeq SBS v3 reagents or on a NextSeq500 high output 75 base pair single-read flow cell using v2 reagents (Illumina, San Diego, CA).

### Data analysis

Adaptors were removed from sequencing read files from miRNA fractions using the clip_adaptors.pl module of mirdeep2^19^. All sequence data were subjected to quality control by fastx_trimmer (http://hannonlab.cshl.edu/fastx_toolkit/commandline.html). Sequence read files from mRNA fractions were trimmed down to 50 bp to remove the poor-quality sequences at the ends. Sequence read files were further collapsed based on sequence identity and annotated for miRNA by mapping to miRbase version 21^20^ using the miraligner module of seqbuster. Reads passing QC standards were mapped to the GRCm38 mouse reference genome assembly using TopHat v2.0.13 (Bowtie v2.0.6) with supplied annotations at parameters (--min-anchor 8, --min-isoform-fraction 0.15). The raw count of mapped reads to genes was determined using the htseq-count utility (http://www-huber.embl.de/users/anders/HTSeq/doc/count.html. Isomirs were identified using the R isomiRs library of the seqbuster package. Differential expression between groups of genes or miRNAs was assessed using DESEQ2^21,22^.

We determined the genomic locations of all mapped mRNA reads and recorded the type of genome region to which the MRE mapped: 3’UTR, 5’UTR, Exon, Intron, Intergenic, or promotor transcriptional start site (TSS), using homer software^21,22^. We then calculated the total read count, as count per million, for each genomic region, normalized by the length of each feature in GRCm38 reference genome. We also performed a similar analysis to determine the chromosomal location of every mRNA read.

Differentially expressed miRNA and mRNA genes were used to perform the Interaction analysis. We selected three comparisons: Early-stage-Normal (EN) vs. Late-State-Tumor (LT), Early-Stage-Tumor (ET) vs. Late-Stage-Tumor (LT), and Late-Stage-Normal (LN) vs. Late-Stage-Tumor that show differential expression of both gene and miRNA features. The rlog transformed data matrices were used to identify the negative correlation using the function cor test from stats package in R. As we were interested only in negative correlations; therefore, we used the alternative hypothesis is “less” to obtain negative correlations between miRNA and genes. The full list of associations obtained from this analysis is shown in Supplemental Table 3. The pairs with correlation < −0.5 and p-value<0.05 are considered strongly correlated in individual comparisons.

### Tumor cell lines and RNA samples

Tumor cell lines 3465KPCA and 3465KPCB were created from lung tumor tissue harvested from a male KP mouse at 16 weeks after intranasal administration of 5×10^6^ PFU of adenovirusCre (Viral Vector Core, University of Iowa, Iowa City, IA). Following euthanasia by cervical dislocation, according to JAX IACUC protocol #01011, lungs were perfused with 10 mls of cold, sterile PBS. Tumors were excised, dissociated with an equal mixture of 4 mg/ml of collagenase D and dispase II (Roche Applied Science, Indianapolis, IN) and cultured in F-12K (Corning, Tewksbury, MA) plus 10% Cosmic calf serum (Hyclone Cytiva Logan, UT) at 37°C and 5%CO2 for until outgrowths were established. Once established, cells were passaged using TrypLE express to dissociate cells (ThermoFisher Scientific Waltham, MA).

3465KPCA and 3465KPCB cells were plated in 6 well dishes transfected with 25-50 pmols mmu-miR-130a-3p, control antimiR (Dharmacon Horizon Discovery, Lafayette, CO) or mCherry mRNA (TriLink Biotechnologies San Diego, CA), as indicated, using Lipofectamine RNAiMAX (ThermoFisher Waltham, MA) following manufacturer’s protocol. Three days after transfection, RNA was isolated from 3465KPCA and 3465KPCB cells using Trizol (Invitrogen ThermoFisher Scientific Waltham, MA) following manufacturer’s protocol for adherent cells. RNA was also isolated from the testes and lungs of an uninfected male KP mouse and from a homozygous male Prdm9 knockout mouse using Trizol and following the manufacturer’s protocol for solid tissues. All RNA samples were quantitated using a Nanodrop1000 spectrophotometer (ThermoFisher Waltham, MA) and 2 micrograms from each sample were DNAse I treated (Promega, Madison, WI) following manufacturer’s protocol. Resulting RNA samples were then reverse transcribed into cDNA using the primer Prdm9intron9 and SuperScript IV (Life Technologies), following manufacturer’s protocol, including the optional RNase H step.

Prdm9exon9: 5’-CCAGGTTGTGAGCTTCTGG-3’

### PCR

RNA levels were determined by PCR analysis of cDNAs. Each PCR reaction contained 100 nanograms of cDNA, 1M betaine, 1X high fidelity PCR buffer, 1 mM of each dNTP, 0.5 micromolar each of primers Prdm9exon5F and Prdm9exon6R, 0.4 units of Phusion High Fidelity DNA polymerase (New England Biolabs Beverly, MA) in a final volume of 20 microliters. Reactions were amplified on an Eppendorf MasterCycler by heating to 98°C for 1 minute, followed by 31 cycles of 98°C for 10 seconds, 68°C for 15 seconds and 72°C for 2 minutes, then 72°C for 5 minutes and hold at 10°C. PCR samples were analyzed on 2% agarose gels (Lonza Rockland, ME), 1X TAE, 0.4 micrograms/ml ethidium bromide and run in 1X TAE.

Prdm9exon5F: 5’-GTGGCTCAGAACACGTCCAG-3’

Prdm9exon6R: 5’-CTTTCTCTCTCGCAGCCTGTAC-3’

#### Sequencing Analysis

Sanger sequencing analysis of *Prdm9* PCR products, generated using primers Prdm9exon5F and Prdm9exon6R, was performed by Eurofins (Louisville, KY).

#### Western blots

Protein extracts were made from 3465KPCA and 3465KPCB cell lines as well as from testes and lung tissues from an uninfected male KP and a homozygous male Prdm9 knockout mouse. Animals were humanely euthanized by cervical dislocation (IACUC protocol #01011), perfused with 10 mls of 1X PBS and desired organs were removed. Excised organs or cultured cells were rinsed three times with cold 1X PBS. Tissues were transferred to ice cold RIPA buffer (Millipore-Sigma Burlington, MA) plus cOmplete mini protease inhibitors (Millipore-Sigma Burlington, MA) and homogenized on ice for 10 seconds using individual sterile pestles (Thermo-Fisher Waltham, MA) with 5 minutes of rest on ice before repeating homogenization. Samples were incubated on ice for 30 minutes with a brief vortex, followed by centrifugation at 10,000 x g for 10 minutes at 4°C. Concentrations of the supernatants was determined by Bradford Assay (Bio-Rad, Hercules, CA) using BSA (Bio-Rad Hercules, CA) for the standard curve. Following denaturation at 95°C for 5 minutes, 60 micrograms of total protein per tissue or 20 micrograms total protein per cell were run on 4-12% Bis-Tris Protein Gel (Thermo-Fisher Waltham, MA) with MOPS SDS Running Buffer (Thermo-Fisher Waltham, MA) and transferred to nitrocellulose (Thermo-Fisher Waltham, MA) using an iBlot Gel Transfer Device (Thermo-Fisher Waltham, MA). The blots were blocked in Bløk-CH Noise Cancelling Reagents for Chemiluminescence Detection (Thermo-Fisher Waltham, MA) at room temperature for 1 hour then incubated with anti-PRDM9 polyclonal antibody^23^ diluted 1:1,000 in Bløk-CH Noise Cancelling Reagent and agitated at 4°C overnight. The next morning the blot was washed with TBST first for 15 minutes, then three times for 5 minutes each then placed in Donkey Anti-Guinea Pig IgG Antibody-HRP Conjugate (Millipore-Sigma Burlington, MA) diluted 1:10,000 in Bløk-CH Noise Cancelling Reagent for 1h at room temperature. After repeating washes with TBST, the blot was developed with SuperSignal West Pico PLUS Chemiluminescent Substrate (Thermo-Fisher Waltham, MA) and imaged with film.

## ACKNOWLEDGEMENTS

We thank Dr. Kenneth Paigen (The Jackson Laboratory) for the generous gift of the anti-PRDM9 antibody and tissues from the *Prdm9* knockout mouse, Dr. Kwok-Kin Wong (NYU Langone Health) and members of his laboratory for technical advice with intranasal infection of mice, Benjamin Low (The Jackson Laboratory) for technical advice with RTPCR, Drs. Olga Anczukow and Hui Tian (The Jackson Laboratory) for helpful discussions and Dr. Michael Wiles (The Jackson Laboratory) for critical reading of this manuscript. We also thank Genome Technologies Sequencing, Genome Engineering Technologies and Protein Sciences cores at The Jackson Laboratory for their expertise and JAX Creative for their assistance with preparing figures. This work was supported in part by NCI (5R21CA155164-2 to J.W. and C.J.B.), the Maine Cancer Foundation (MCF JMW-01 to J.W. and C.J.B.) and the Knowlton Family Foundation (to C.J.B.). Research reported in this publication was partially supported by the National Cancer Institute of the National Institutes of Health under Award Number P30CA034196.

## AUTHOR CONTRIBUTIONS

J.W. and R.S.M. designed experiments and JW performed all RIP-SIR and RTPCR experiments and generated all high-throughput sequencing libraries. R.S.M. generated lung tumor cell lines. A.S., R.K. and K.S. analyzed all high-throughput sequencing data. S.L.D., R.P.L. and H.J.M performed all high-throughput sequencing and R.D. examined and diagnosed all histology samples. R.L.S performed the western blot experiment. J.W. wrote the manuscript with input from other authors.

