## Supplemental Figs 1, 2 for "RNA Immunoprecipitation and Sequencing of Isolated RNAs (RIP-SIR) Identifies Endogenous miRNA-Target Interactions"

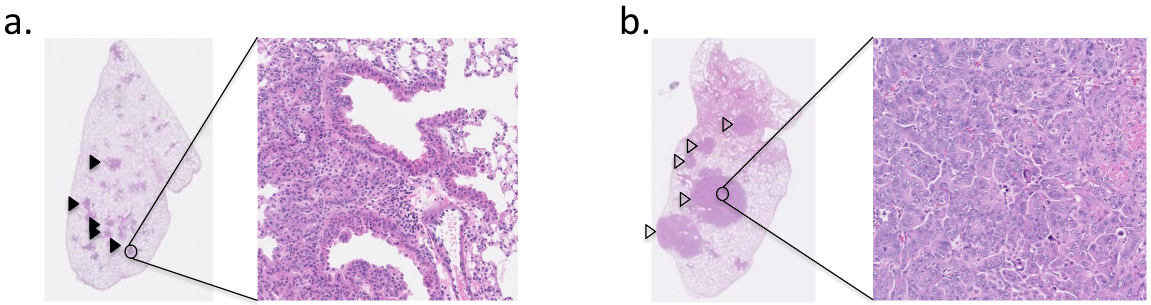


**Supplemental Fig.1 Histology of K and KP mouse lung tumors**. **a**, 16 weeks after infection with adenovirus Cre, hemotoxylin and eosin staining of the left lung lobe shows a large number of lesions (solid arrowheads) that are primarily adenomas. One of these lesions is enlarged in the panel on the right. **b**, In contrast, the left lung lobe of a KP animal at 16 weeks after infection shows a large number of lesions (empty arrowheads) that are predominantly adenocarcinomas. One of these lesions in enlarged in the panel on the right.

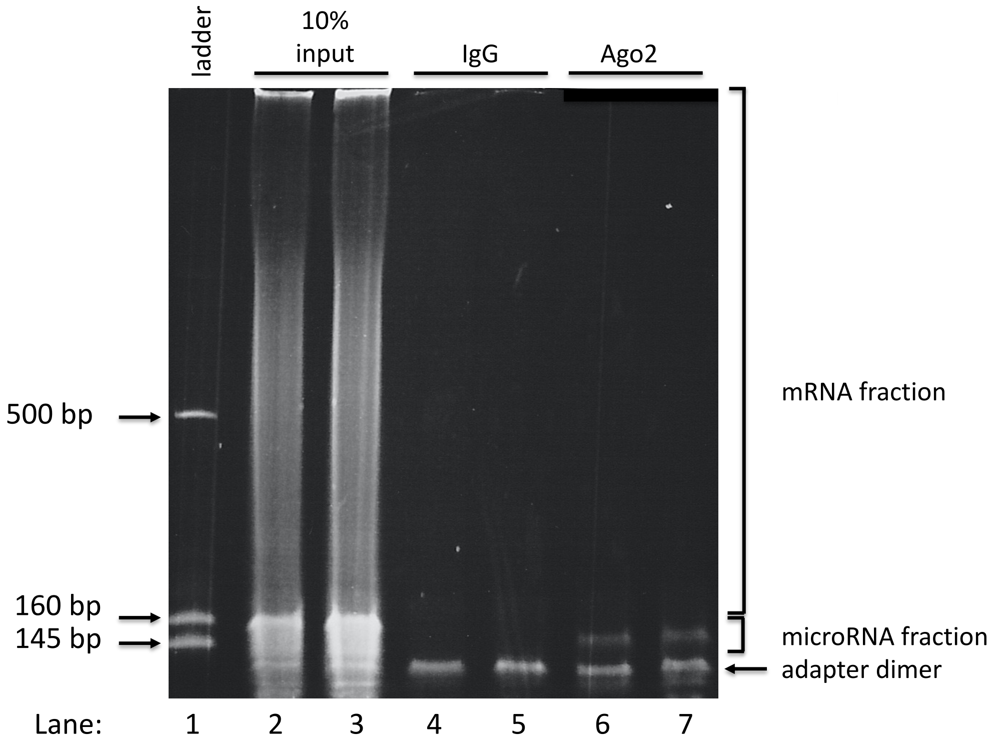

**Supplemental Fig. 2 Size fractionation of RIP-seq libraries**. RIP-seq libraries were generated following immunoprecipitation of crosslinked whole mouse lung tissues with an antibody against either Ago2 (lanes 6, 7) or an IgG isotype control (lanes 4,5). As a positive control, a sequencing library was also generated from 10% of the input crosslinked mouse lung extract that was not immunoprecipitated (lanes 2, 3). Lane 1 contains an RNA ladder to indicate the sizes of the libraries. All libraries were run on a 5% acrylamide gel and stained with ethidium bromide. Libraries between 140-170 nts were labeled as the miRNA fractions while libraries between 170-500 nts were labeled as the mRNA fractions. The band corresponding to the adapter dimer is also indicated.
