## Supplemental Table 4 for "RNA Immunoprecipitation and Sequencing of Isolated RNAs (RIP-SIR) Identifies Endogenous miRNA-Target Interactions"

**Supplemental Table 4** Locations of identified MREs in analyzed transcripts

| Gene | RIP-seq read # | Transcript location | Genomic location | Distance | Mapped location | Transcript length (nts) | # of exons | 5'UTR (nts) | 3'UTR (nts) | Coding (nts) |
|---|---|---|---|---|---|---|---|---|---|---|
| Hmga2 | 1 | exon 6 | 120361603 | | MRE1 | 115,661 | 6 | 872 | 3,014 | 111,775 |
| | 2 | exon 6 | 120361607 | 4 | MRE1 | | | | | |
| SIR probe | 2 | | 120362825 | | | | | | | |
| Notch 2 | 1 | intron 1 | 98045911 | | MRE 1 | 136,830 | 34 | 161 | 2,973 | 133,746 |
| | 2 | intron 1 | 98046027 | 116 | MRE 1 | | | | | |
| | 3 | intron 5 | 98088301 | 42,274 | MRE 2 | | | | | |
| | 4 | intron 5 | 98088314 | 113 | MRE 2 | | | | | |
| | 3 | intron 17 | 98118519 | 30,205 | MRE 3 | | | | | |
| | 6 | intron 17 | 98118545 | 26 | MRE 3 | | | | | |
| SIR probe 1 | | | 98118454 | | | | | | | |
| SIR probe 2 | | | 98045841 | | | | | | | |
| Prdm9 | 1 | intron 9 | 15549547 | | MRE1 | 20,253 | 10 | 894 | 30 | 19,329 |
| | 2 | intron 5 | 15556423 | 6,876 | MRE2 | | | | | |
| | 3 | intron 5 | 15556450 | 27 | MRE2 | | | | | |
| SIR probe | | | 15556360 | | | | | | | |
| Ppp1r16b | 1 | intron 1 | 158672694 | | MRE1 | 99,705 | 11 | 29,331 | 4,471 | 65,903 |
| | 2 | intron 1 | 158672743 | 49 | MRE1 | | | | | |
| | 3 | intron 1 | 158681961 | 9,218 | MRE2 | | | | | |
| | 4 | intron 1 | 158681991 | 30 | MRE2 | | | | | |
| | 5 | intron 1 | 158689973 | 7,982 | MRE3 | | | | | |
| | 6 | intron 1 | 158690036 | 160 | MRE3 | | | | | |
| | 7 | intron 1 | 158690127 | 91 | MRE3 | | | | | |
| | 8 | intron 1 | 158690128 | 1 | MRE3 | | | | | |
| | 9 | intron 1 | 158690133 | 5 | MRE3 | | | | | |
| | 10 | intron 2 | 158713429 | 23,296 | MRE4 | | | | | |
| | 11 | intron 2 | 158713375 | 46 | MRE4 | | | | | |
| | 12 | exon 11 | 158761361 | 47,986 | MRE5 | | | | | |
| | 13 | exon 11 | 158761496 | 135 | MRE5 | | | | | |
| SIR probe | | | 158690036 | | | | | | | |
